# Toward computational design of protein crystals with improved resolution

**DOI:** 10.1101/657262

**Authors:** Jeliazko R. Jeliazkov, Aaron C. Robinson, Bertrand García-Moreno E., James M. Berger, Jeffrey J. Gray

## Abstract

Substantial advances have been made in the computational design of protein interfaces over the last 20 years. However, the interfaces targeted by design have typically been stable and high affinity. Here, we report the development of a generic computational design method to stabilize the weak interactions at crystallographic interfaces. Initially, we analyzed structures reported in the Protein Data Bank (PDB) to determine whether crystals with more stable interfaces result in higher resolution structures. We found that, for twenty-two variants of a single protein crystallized by a single individual, Rosetta score correlates with resolution. We next developed and tested a computational design protocol, seeking to identify point mutations that would improve resolution on a highly stable variant of staphylococcal nuclease (SNase Δ+PHS). Only one of eleven initial designs crystallized, forcing us to re-evaluate our strategy and base our designs on an ensemble of protein backbones. Using this strategy, four of the five designed proteins crystallized. Collecting diffraction data for multiple crystals per design and solving crystal structures, we found that designed crystals improved resolution modestly and in unpredictable ways, including altering crystal space group. *Post-hoc*, *in silico* analysis showed that crystal space groups could have been predicted for four of six variants (including WT), but that resolution did not correlate with interface stability, as it did in the preliminary results. Our results show that single point mutations can have significant effects on crystal resolution and space group, and that it is possible to computationally identify such mutations, suggesting a potential design strategy to generate high-resolution protein crystals from poorly diffracting ones.

## 1 Introduction

X-ray crystallography is still the primary method for acquiring atomic-resolution structural information about biological macromolecules such as proteins, and it is indispensable for gaining functional and mechanistic insights across biological and pharmacological disciplines^1^. However, because of its highly unpredictable nature, crystallography is viewed more as art than as science—a fact reflected by the low rate of success in large-scale protein crystallization efforts (~10–20%)^2,3^. Even when proteins produce diffraction-quality crystals, the data may be of low quality or difficult to solve. Approximately 23% of the crystal structures reported to the PDB diffract to a resolution of 2.5 Å or worse^4^. At a resolution of 2.5 Å, the backbone, side chains, and small molecules can be fit with a reasonable degree of precision to the electron density; however, key features such as the placement of water molecules or alternate side-chain conformations may be less certain. At even lower resolutions (3–6 Å), ligands/side chains and even the main chain may not be fit reliably^5–8^. An inability to resolve ligands, water molecules, small molecules, or side-chain interactions prevents accurate understanding of catalytic mechanisms, drug-protein interactions, or the organization of certain macromolecular complexes, and precludes computational design from using natural proteins as input.

Historically, rational design has been used to overcome various degrees of protein recalcitrance to crystallization, from improving existing crystals to generating new ones^9^. The variety in strategies has been quite broad. Some strategies can be applied when only the protein sequence is known, even before crystal trays are laid; *e.g.*, by deleting loops or regions of low sequence complexity^10^, or by identifying stabilizing mutations from homologous sequences^11^, surface entropy reduction (SER)^12^, or *de novo* crystal design^13^. Other strategies have focused on improving an existing crystal, such as through the rational engineering of crystal contacts^14,15^. Of the above strategies, all but SER must be tailored to a specific target protein. The necessity for protein-specific approaches is somewhat surprising considering that the underlying physics of crystallization is universal. For example, Fusco *et al.* identified two generic mechanisms underlying crystal formation in their analysis of 182 proteins in 1,536 crystallization conditions^16^. In principle, a reliable and general method for enhancing the resolution of poorly-diffracting crystals through rational and computational design should exist.

Here, we report our attempts to develop resolution-enhancing computational design of protein–protein interactions at crystallographic interfaces. We began by identifying physical determinants of high-resolution protein crystals. This led us to identify a positive correlation between resolution and crystal lattice ‘stability’ (the Rosetta-determined score of the asymmetric unit and the unique protein–protein interactions defining the crystal lattice). A Rosetta protocol was then developed to identify stabilizing (resolution-enhancing) point mutations. This protocol was benchmarked *in silico* against rationally-engineered protein crystals^17^. We tested our protocol experimentally by designing, cloning, expressing, purifying, and crystallizing variants of a highly stable form of staphylococcal nuclease (SNase). We found that variants designed on a single, fixed backbone crystallized rarely (1/11), whereas variants designed on an ensemble of backbones crystallized more readily (4/7). Comparison of the highest-resolution shells for collected diffraction data (determined by CC_1/2_) revealed only minor improvements in resolution (~0.05 Å) for three of five designs that crystallized. Surprisingly, two of the resolution-enhancing designs altered the crystal space group. An analysis of our efforts shows that space group changes could have been predicted for three of the five designs, but that crystal lattice stability does not correlate with resolution for our test protein.

## 2 Materials and Methods

All data associated with this paper are available online via Zenodo at the following DOIs:

- 10.5281/zenodo.3216968
- 10.5281/zenodo.3222946
- 10.5281/zenodo.3228344
- 10.5281/zenodo.3228838
- 10.5281/zenodo.3235486
- 10.5281/zenodo.3235518
- 10.5281/zenodo.3228351

### 2.1 Curation of Crystal Datasets

Three protein structure datasets were constructed on October 10^th^, 2016, from the Protein Data Bank^4^ (PDB), termed the: “PDB representative”, “SNase”, and “Mizutani” sets.

To generate the PDB representative set, we first generated three lists of PDB IDs and then took the intersection of the lists. The first list ensured that we only analyzed non-redundant, reasonable-quality protein structures. Using the PISCES server^18^, we generated a list of non-Cα-only X-ray structures in the PDB adhering to the following criteria (culling by chain):

- 25% maximum sequence identity,
- resolution better than 3.0 Å,
- R-value < 0.3, and
- Proteins comprising more than 40, but less than 10,000 residues.

This list, `pisces.txt`, contained 10,886 PDB IDs. Next, we generated a second list to limit our analysis to solely crystallographic protein–protein interactions. Using the advanced search option on the PDB website (https://www.rcsb.org/pdb/search/advSearch.do), we generated a list of PDB structures with monomeric stoichiometry and only a single chain in both the biological and asymmetric units. This list, `pdb.txt`, contained 36,899 PDB IDs. Finally, we generated a third list to exclude structures containing many ligands or non-protein atoms. Starting with the PDB IDs from the pisces.txt list, we used a Python script (`1-parse-pdb-remarks.py`) to filter PDB IDs, selecting for the absence of REMARK 465/470/475/480 records, which indicate missing atoms, and a fraction of non-HOH HETATM records greater than 0.1. This list, `missing_or_nonhet.txt`, contained 873 PDB IDs. We took the intersection of our three lists of PDB IDs as our “PDB representative” set; this was performed in R using the `merge-three-lists.R` script. The final list contained 379 PDB IDs.

Separately, to generate the SNase set, we used a new PDB advanced search to select for X-ray structures with UniProt Accession ID: P00644, in addition to the above criteria for stoichiometry, biological unit, and asymmetric unit, but not culling for sequence identity, R-value less than 0.3, absence of atoms or presence of ligands. The SNase set contained 256 PDB IDs, which were not present in the PDB representative set.

Finally, the Mizutani set was simply composed of the 21 structures of diphtine synthase deposited by Mizutani *et al.* in their study of rational crystal contact engineering^17^. These structures were not present in either the PDB representative or SNase set.

### 2.2 Modeling of Crystals

In this study, we sought to computationally quantify crystallographic protein–protein interactions. To that end, proteins were modeled in three states: (1) as a crystal, including all symmetry mates within 12 Å, (2) as a collection of pairwise interfaces, and (3) as a monomer. These states were constructed for each dataset. Furthermore, pairwise interfaces were analyzed with the Evolutionary Protein–Protein Interface Classifier^19^ (EPPIC) to verify that the interactions we were assessing were crystallographic and not biological.

#### PDB Representative

Monomers were downloaded from the PDB and energy minimized using Rosetta, weekly version: v2018.24. To ensure accurate energy calculation and because Rosetta cannot model all ligands/co-factors/*etc.*, HETATM records were omitted. Energy minimization was performed by the FastRelax protocol^20,21^ with the following command line:

~~~
relax.linuxgccrelease -l
list.txt-relax:constrain_relax_to_start_coords -relax:ramp_constraints -ex
1 -ex2 -use_input_sc –flip_NHQ -no_optH false -nstruct 10 -out:pdb_gz,
~~~

where `list.txt` contained the PDBs. Individual crystallographic interfaces were generated and analyzed using the pre-compiled EPPIC command-line interface (version 3.0.5):

~~~
epicc -i 1ABC.pdb -l -p -s,
~~~

where 1ABC.pdb is any PDB. Any PDB ID with an interface predicted by EPPIC to be biological and not crystallographic was excluded from further analysis if the corresponding PDB entry or supporting literature indicated that the biologically relevant state was not monomeric (as we only wish to study crystallographic interactions). Crystals were modeled using the same protocol, except this time the `“-symmetry:symmetry_definition CRYST1”` flag was included to enable modeling of the asymmetric unit and all symmetry mates within 12 Å as previously described^22^.

The energy of crystallization was computationally determined using the weekly PyRosetta^23^ release, v2018.24, and the November 2016 version of the Rosetta scoring function^24,25^. The script `score_crystal_interfaces_parallel.py` evaluated the energy of each crystallographic interface, which was later combined with the energy of the monomer to yield the crystal energy (see Results 3.1). After excluding four structures that could not be modeled with our approach and twelve structures that accidentally included biological interactions in the crystal, 364 PDBs were analyzed from the PDB. These PDBs are listed in: `pdb-representative-rosetta.txt`.

#### SNase

As above, SNase monomers were downloaded from the PDB and energy minimized using Rosetta v2018.24-dev60250. Unlike above, HETATM records were retained, because the nucleotide analog thymidine-3’,5’-diphosphate (THP) bound by the enzyme makes important crystal contacts. The ligand geometry was fixed in simulations and read from the PDB Chemical Components Dictionary. Energy minimization was performed as described above. Individual crystallographic interfaces were not generated and analyzed using EPPIC because SNase is known to be a monomer. Crystals were modeled and crystal energies were computed as described above.

#### Mizutani

Modeling of the 21 diphtine synthase structures from Mizutani *et al.* was performed similarly to modeling of the PDB and SNase crystal structures, as described above. The only exception being that diphtine synthase is naturally a dimer, so during energy evaluation the two dimeric chains were treated as a monomer and only crystallographic, not biological, interfaces were considered. As with SNase, because the biological state is known, EPPIC was not used to generate or analyze crystallographic interfaces. Crystals were modeled and crystal energies were computed as described above.

### 2.3 Forward Design of Diphthine Synthase

To determine the computational design approach most likely to be experimentally successful, we tested several strategies on diphthine synthase. Since the resolution is reported for twenty variants, we can ask if a particular design strategy can predict resolution-improving variants. This approach is known as forward design. The energy-minimized structure of wild-type diphthine synthase (PDB ID: 1WNG) was used as input for design. At each position mutated by Mizutani *et al.* (26, 49, 54, 65, 69, 79, 140, 142, 146, 171, 173, 187, and 261), each amino acid except cysteine or proline was tested. We attempted to accommodate the mutation by permitting varying degrees of freedom in the wild-type crystal form (simulated by Rosetta Symmetry). These different design strategies ranged from only permitting neighboring side chains to repack to re-docking in the crystal lattice (see Results). All energy calculations were performed in the crystal form and averaged over ten repeats.

### 2.4 Computational Design of SNase

Design for SNase variants initially followed our most successful diphthine synthase approach: introducing a point mutation at a position followed by repacking of side-chains followed by energy minimization of side-chain and backbone dihedral angles. An energy-minimized structure of Δ+PHS SNase (PDB ID: 3BDC) was used as input. The geometry of the THP ligand was held fixed during the simulation and defined by a Rosetta params file derived from the PDB coordinates. Later, we introduced an ensemble of 200 perturbed backbones generated by Rosetta Backrub^26^ as it has been shown that interface ΔΔG prediction is more accurate when using an ensemble of backbones rather than just a single input^27^. The following steps were repeated for each member of the backbone ensemble and for every designable surface position, defined as residues having a Cα–Cα distance under 8 Å across any crystallographic interface. First, the position was mutated to one of eighteen amino acids (cysteine and proline were excluded). Then, the crystal form was generated using Rosetta Symmetry in the wild-type space group and unit cell dimensions. Finally, side-chains were repacked to accommodate the mutation in the crystal form. The error in this modeling was calculated across the ensemble of 200 backbones, instead of by repeating the simulation ten times.

### 2.5 Cloning, Expression, and Purification of Proteins

Point mutations were introduced by Quikchange mutagenesis^28^ into the highly stable Δ+PHS variant^29^, expressed in *E. coli* BL21/DE3 cells transformed with the pET-24a+, and purified as previously described^30^.

### 2.6 Protein Crystallization

Crystals of Δ+PHS and its variants were grown by the hanging drop vapor-diffusion method at 277 K. The reservoir solution varied, ranging in pH from 6–9, with 20–40% (v/v) 2-methyl-2,4-pentanediol (MPD), either 3 or 2 molar equivalents of THP, either 2 or 1 molar equivalents CaCl_2_, and 25 mM potassium phosphate. The protein concentration varied across variants, but was always mixed in a 1:1 ratio with the reservoir solution to make the drop. Conditions are detailed for each crystal in Supplemental Table 1. Crystals typically appeared after one week, were harvested with Hampton Research CryoLoops™ on CrystalCap™ Copper HT magnetic sample mounts, and were immediately flash-cooled in liquid nitrogen. Crystals were stored at 77 K until data collection.

### 2.7 Data Collection and Structure Determination

All X-ray diffraction data were collected for single crystals at 77 K using a Rigaku FR-E SuperBright rotating anode X-ray generator and a Rigaku DECTRIS PILATUS 200K pixel array detector. Diffraction data were indexed, integrated, and scaled using the XDS program package^31^. Phasing, modeling building and model refining were performed using PHENIX^32^. Phasing was performed by molecular replacement in PHASER^33^ using the search model 3BDC, with solvent-exposed or mutated side chains truncated to the Cα position to avoid biasing side-chain placement at crystal contact sites.

Side chains were rebuilt using COOT^34^: initial placement was performed using the Mutate and Autofit function and followed by manual refinement. Whole-structure refinement and water placement were performed using *phenix.refine*^35^ and *phenix.rosetta_refine*^36^. Data collection and refinement statistics are shown in Supplemental Table 2. Crystal structures deposited to the PDB include models 6OK8 (K127L), 6OK9 (K133M), and 6OKA (Q123D).

## 3 Results

Our primary goal in this effort was to develop a broadly applicable, Rosetta-based computational method for crystal contact design, with the goal of systematically predicting single point mutations that could enhance resolution. To this end, we first asked whether or not Rosetta scoring functions might correlate with resolution.

### 3.1 Rosetta Score Correlates with Resolution, When Other Variables Are Controlled

With over 129,000 crystal structures of biological macromolecules^4^, the PDB provides a trove of data that can be used to determine whether or not Rosetta score correlates with the resolution of protein crystals. To ensure a fair comparison, structures must be first energy-minimized in the Rosetta scoring function using the Rosetta FastRelax protocol^20,21^. As FastRelax runtime scales with protein size, testing every structure in the PDB is not feasible. Furthermore, some structures are overrepresented in the PDB, which might bias analysis. Instead of analyzing all structures, we selected a diverse and representative subset of the PDB containing 364 structures, which had a maximum sequence identity of 25%, a resolution better than 3 Å, R-values less than 0.3, and featured only crystallographic interactions (fully described in Methods 2.1 & 2.2). For every structure in the set, we generated all symmetry-mates within 12 Å using Rosetta Symmetry^22^ and energy minimized this “crystal” form ten separate times using the default FastRelax protocol, which features four cycles of minimization each with progressively weaker harmonic constraints to prevent substantial deviation from the starting coordinates. Separately, we energy-minimized the monomeric form of the protein, which was also the asymmetric unit and the biologically-relevant unit. We approximated the energy of the crystal as 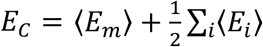, where 〈*E*_*m*_〉 is the average Rosetta score of ten energy-minimized monomers and ∑_*i*_〈*E*_*i*_〉 is the average Rosetta score of interface *i* in the crystal form, summed over all interfaces within 12 Å of the asymmetric unit. Hence, *E*_*c*_ represents that energy of the minimal unit required to generate the crystal. The relationship between resolution and score is shown in Figure 1A.

**Figure 1:**
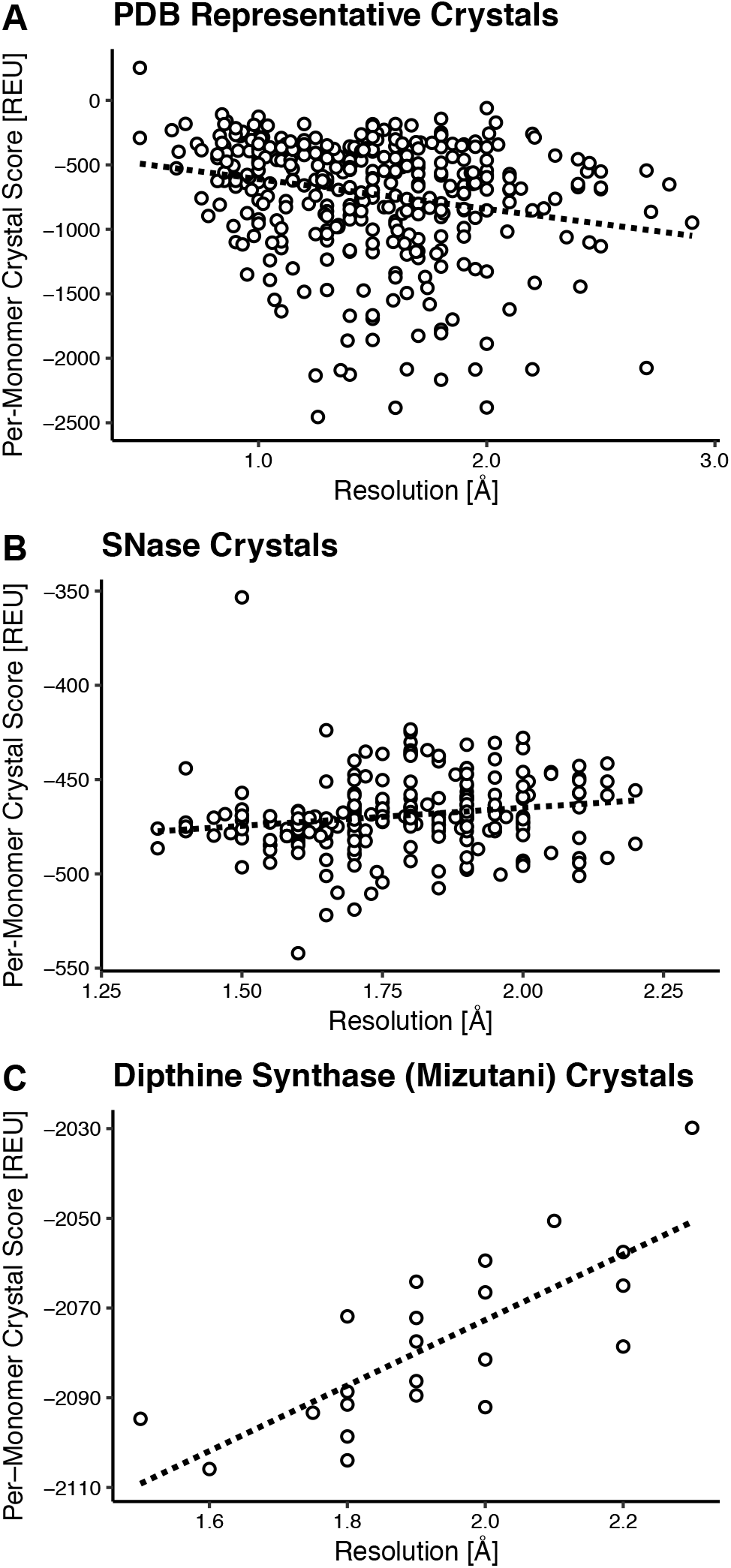
Rosetta score correlates with resolution when comparing crystals of the same protein varying only by point mutations. The per-monomer crystal scores (*i.e.* the score of the monomer plus all crystallographic interactions, which is the minimal interacting unit required to generate the crystal) and crystal resolutions are compared for three sets of protein structures. The PDB-representative set (A) samples 364 monomers with distinct sequences and shows no relationship between score and resolution. The SNase set (B, excluding an outlier at 2.5 Å) compares only crystals of *Staphylococcal aureus* nuclease, attempting to rule out the protein as a variable, but still there is no trend. Finally, the dipthine synthase set (C) compares crystals that vary only by a point mutation, ruling out most extrinsic variables, and score correlates with resolution.

We initially found a slight anti-correlation between resolution and score for the representative PDB set: higher resolution structures had lower scores. We hypothesized that this unexpected result was caused by our inability to control for the many variables that affect resolution that are not captured by Rosetta score (*e.g.*, how the highest resolution-shell cutoff was decided, user handling of the crystal, the content of the reservoir solution, *etc.*). To test this hypothesis, we analyzed two additional sets of crystal structures in the same manner as the PDB representative set. The first additional set we analyzed controlled for the protein as a variable. We searched for a small, globular protein, with many structures in the PDB that differed only slightly from each other, but that spanned at least 1 Å in resolution. Of the multiple proteins fulfilling these criteria, we selected SNase, which had 256 crystal structures. We used Rosetta Symmetry and FastRelax to generate ten energy-minimized monomers and crystals, and computed the energy of the crystal as described above. The SNase set did not show a strong correlation between resolution and score (Figure 1B). To further control extrinsic variables, we analyzed twenty-two variants of a single protein (diphthine synthase) that had been previously cloned, expressed, purified, crystallized and the crystal structures solved by one scientist in a crystal engineering study by Mizutani *et al.*^17^. Despite the variants only differing by a point mutation or two, the crystal structures spanned a range of resolutions from 1.5 Å to 2.3 Å. The structures were energy minimized and the crystal energies calculated in a similar fashion as for the previous two sets. The Mizutani set showed a strong correlation (R^2^ = 0.8) between Rosetta score and resolution (Figure 1C).

### 3.2 Rosetta Can Identify Resolution-Enhancing Mutations

Since low score corresponded to high resolution for the Mizutani set, we next sought a fast computational design strategy that could identify resolution-enhancing mutations from the wildtype (WT) crystal structure. We tested six strategies on the Mizutani set in an approach known as forward design. Using the energy-minimized crystal form of the wildtype protein as input, we introduced point mutations one-by-one at the positions engineered by Mizutani *et al*. We then optimized side-chain dihedral angles while keeping the backbone fixed (side-chain repacking)^37^. Following repacking, we tested six design strategies with varying degrees of freedom: (1) we did nothing else before evaluating the score, (2) we applied gradient-based energy minimization on side-chain dihedral angles, (3) we applied gradient-based energy minimization on side-chain and backbone dihedral angles, (4) we applied gradient-based energy minimization on side-chain dihedral angles and the relative position/orientation of the protein and its symmetry mates, (5) we applied gradient-based energy minimization on side-chain and backbone dihedral angles and the relative the relative position/orientation of the protein and its symmetry mates, and (6) we sampled the relative position/orientation of the protein and its symmetry mates, translating in steps of 0.05 Å and rotating in steps of 0.1 degrees, followed by energy minimization as in Strategy 5 over four Monte Carlo cycles. All gradient-based minimization was run until convergence was achieved, defined as a change in Rosetta score of less than 0.00001 following an iteration of minimization, or for 200 iterations. Each strategy was tested ten times to assess error. The forward design results for all strategies are shown in Supplemental Figure 1.

We found Strategy 3 (minimizing on side-chain and backbone torsion angles after repacking) to be the most successful. Figure 2 compares the predicted change in score between each variant and WT diphthine synthase for Strategy 3. This approach successfully predicts 6 of 17 (35.3%) resolution-enhancing mutations identified in the paper. In addition to these six, this approach predicts 46 other mutations to have lower energy than WT and thus could be potentially resolution-enhancing; however, these were not experimentally characterized by Mizutani *et al*., so it is unclear if these predictions are correct. Assuming a worst-case scenario where these uncharacterized point mutations do not enhance resolution-enhancing mutations, this design approach would predict 6 resolution-enhancing mutations out of 52 possibilities or 11.5%. Both success rates, 35.3% and 11.5%, compare favorably with historical protein interface design success rates, which are typically under 10%^38^.

**Figure 2:**
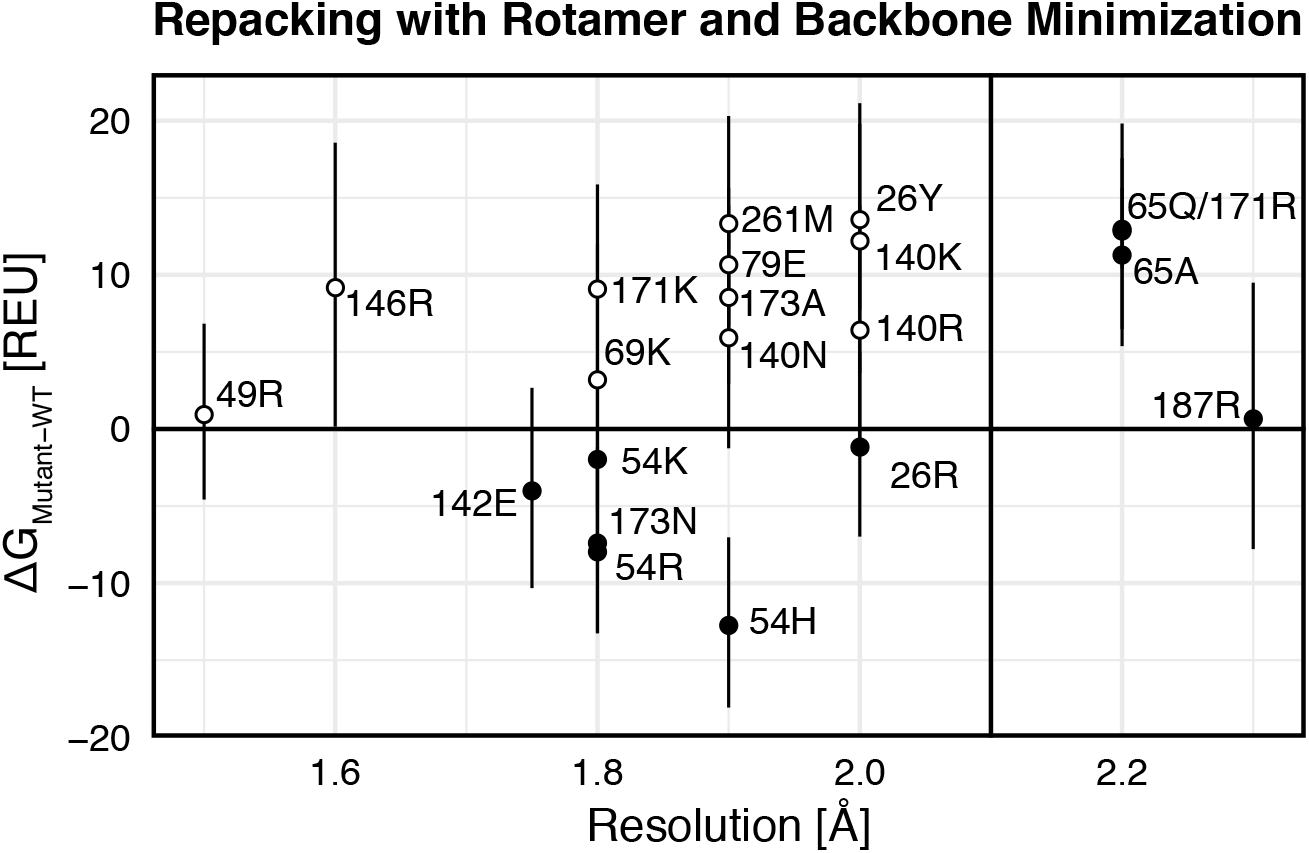
Forward design on dipthine synthase suggests that Rosetta can successfully identify mutations that improve or worsen resolution. The plot shows the difference in score (in the crystal form) between WT and various designs versus the experimentally determined resolution, with the solid lines indicating the WT values. The black points are mutations whose score correctly predicts the sign of the resolution change (*i.e.* better than WT score results in better than WT resolution and *vice versa*) whereas the hollow points are mutations whose score incorrectly predicts resolution change. Standard deviations in score are calculated from 10 repeats of the design simulation.

### 3.3 Rosetta Designed Crystals Do Not Significantly Improve Resolution

To determine whether our design approach was applicable to other proteins, we tested it on a model system: Δ+PHS, a well-characterized highly stable variant of staphylococcal nuclease (SNase)^29^. We identified candidate designable residues at crystallographic interfaces as those with a Cα–Cα distance under 8 Å to neighboring symmetry mates. At each position, we introduced a point mutation followed by side-chain repacking and energy minimization of side-chain and backbone dihedral angles. We selected eleven designs for experimental characterization. However, of the eleven, only a single variant crystallized in conditions where the WT protein normally crystallizes. Since we wanted our approach to yield crystals without having to reoptimize crystal growth conditions, we sought to improve the crystallization rate of our designs. To this end, we introduced a step to generate backbone diversity before design. We drew inspiration from recent work showing that interfacial ΔΔG calculations are more accurate when the change in energy is computed across an ensemble of models^27^. Since backbone diversity was introduced beforehand, we were more conservative in our approach and we followed the introduction of point mutations with only side-chain repacking (Strategy 1 from 3.1). From the second round of design we identified seven possible variants, but since two overlapped with those found in the first round, only the five new variants were characterized experimentally. With this design approach, four of the seven variants yielded crystals in WT-like conditions; thus, designing on an ensemble of structures had improved our crystallization rate from 9% to 57%.

Next, we determined the resolution of diffraction data collected for our variants and compared it to that of the WT protein. To control for differences across crystals, we collected full diffraction data sets for at least three crystals of each variant (up to a maximum of fifteen), depending on the propensity of each variant to form diffraction-quality crystals. The K127L variant, in particular, affected crystal growth and nucleation significantly, yielding larger crystals across more conditions than the other variants. We then indexed, integrated, and scaled the data sets using XDS. The most likely space group was determined by POINTLESS^39^. For all variants except K64R and K127L, it was the WT space group (P2_1_). We found that K64R crystallized in P2_1_2_1_2_1_ and K127L crystallized in P4_1_, the third and second most common space groups for SNase crystals, respectively. In total, we were able to process the data for 37 of the 43 crystal diffraction patterns collected, with the remaining six datasets failing to index due to issues such as ice rings or poor spot profiles. We report a summary of the collected and processed diffraction data in Supplemental Table 1.

Following processing with XDS, we identified the highest resolution shell as the shell with the highest resolution still having a significant CC_1/2_ (*t* < 0.01). We used CC_1/2_, or the correlation between intensities when the data is split in half, to select the highest resolution shell because it provides a rigorous statistical cutoff^40^. To compute significance we calculated a *t*-value, 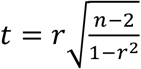, where *r* is the CC_1/2_ value and *n* is the number of reflection pairs (the degrees of freedom), and compared it to Student’s *t*-distribution with the same degrees of freedom^41^. We found that, on average, three of the five designs (60%: Q123D, K64R, and K127L) achieved a higher resolution than WT (Figure 3).

**Figure 3:**
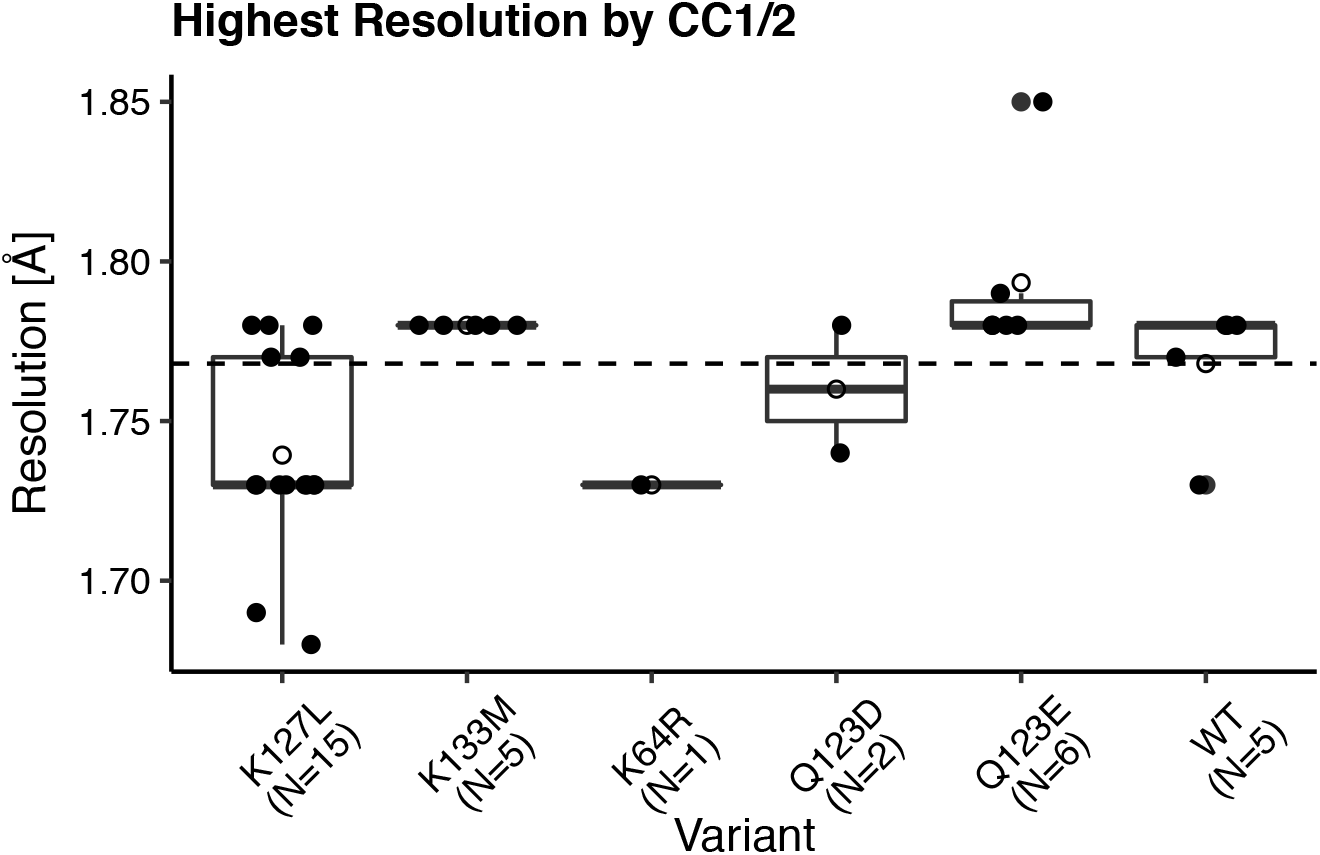
Distributions of the highest resolution shell show that some designs improve on WT resolution. Data were collected from multiple X-ray diffraction experiments and determined by significant CC_1/2_ according to Student’s *t*-test. Boxplots show the median resolution ± one quartile. Open circles indicate the average resolution. The dashed line is the average WT resolution. Designs K127L, K64R, and Q123D have higher average resolution than WT.

However, the improvement was minimal (<0.05 Å). In general, variant resolutions fell within a very narrow range: 1.67–1.85 Å, the width of which was only slightly greater than the range typically spanned by the resolutions of multiple crystals from the same variant (~0.1 Å).

### 3.4 Rosetta-Designed Crystals Do Not Behave as Predicted

Intrigued by the unexpectedly small variation in resolution between designed variants, we solved the crystal structures of several candidates to ask whether there was an underlying structural basis for the small changes in resolution. We discuss the variants below, grouped by their observed effects on SNase crystallization, and provide a general summary of observations across all variants.

#### 3.4.1 Q123D and Q123E

*In silico*, the Q123D design strengthened the crystallographic interface by introducing an electrostatic interaction between D123 in the asymmetric unit and K71 in a neighboring symmetry mate. Upon solving the crystal structure, we found minor changes (less than 0.5 Å RMSD) in the backbone conformation (Supplemental Figure 2). Analysis of the site around the Q123D mutation revealed that residue D123 interacted with residue K84 of a neighboring symmetry mate (with a 3.1 Å distance between the lysine nitrogen and aspartic acid oxygen), instead of K71 (Figure 4). This result is in contrast to the WT structure, where residue K71 interacts with Q123 (3 Å distance between the corresponding oxygen and nitrogen atoms) and K84 solely contacts a phosphate oxygen of THP (Supplemental Figure 3).

**Figure 4:**
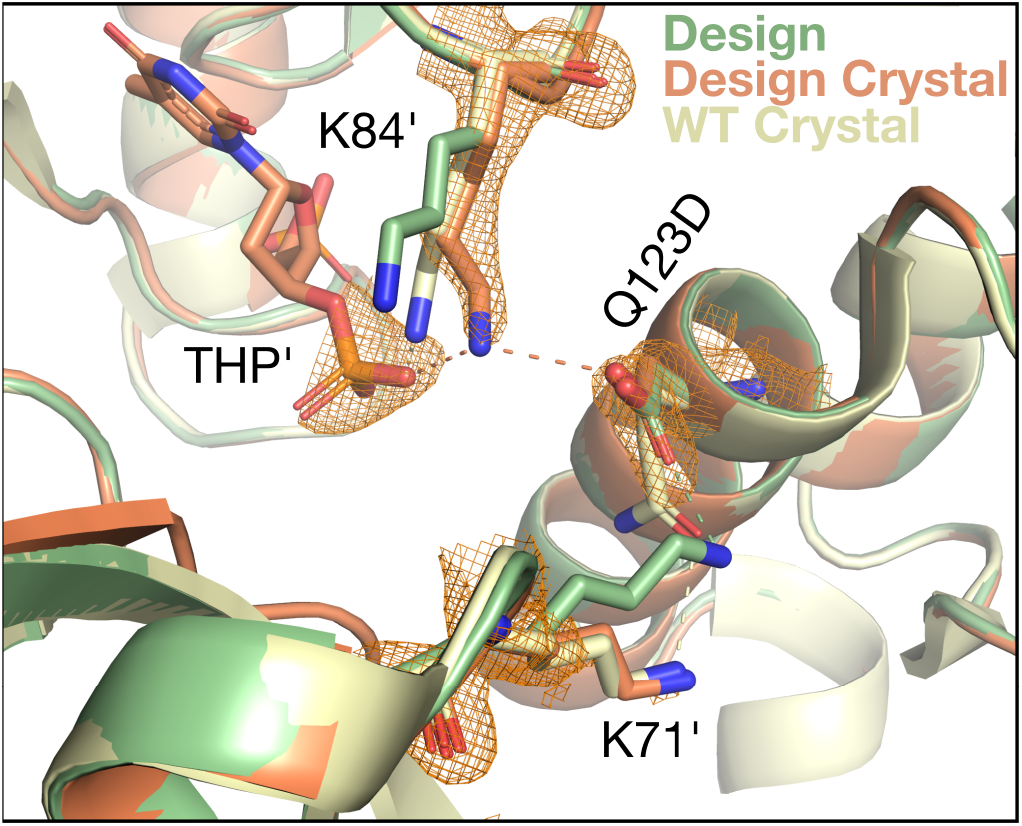
Q123D design (green, PDB ID 6OKA) predicts the correct interaction type, but the incorrect interaction residue. In the design, residue D123 is predicted to form an electrostatic interaction with residue K71’ (‘ indicates symmetry mate), improving on the WT Q–K interaction (pale yellow). However, this interaction is missing in the density and crystal structure of the variant (both orange; the 2mFo-DFc map contoured at 1.5σ for the Q123D variant crystal structure is carved within 2 Å of residues 71, 84, and 123). In place of the Q123–K71 interaction, D123 hydrogen bonds to K84, which also non-covalently interacts with the nucleotide analog (thymidine-3’,5’-diphosphate, THP) bound in the SNase active site. In this figure, each residue belongs to either the asymmetric unit or a different symmetry mate. Key interactions with atom-pair distances under 3.5 Å are shown as dashed lines.

Although we were unable to solve the crystal structure for the Q123E variant due to twinning and the presence of ice, we expect a similar interaction to be occurring. This supposition is supported by the observed average resolutions, which are quite similar to the WT protein: Q123D slightly improves (by 0.01 Å) resolution whereas Q123E slightly worsens resolution (by 0.02 Å).

#### 3.4.2 K133M

Like Q123E and Q123D, the K133M variant did not significantly alter resolution with respect to the WT protein and resulted in minimal backbone movements (0.17 Å RMSD, Supplemental Figure 2). The design was favored *in silico* because it replaced an unfavorable electrostatic interaction between K133 and H8 with a van der Waals contact between M133 and H8, while also slightly reducing the entropic cost of forming that crystal contact (Figure 5). However, the K133M crystal structure revealed that, although the interface had compacted slightly, the side chains were too distant to interact. Compared to the design, the minimum distance between M133 and H8 in the crystal grew from 3.6 Å to 5.3 Å. This lack of interacting side chains likely explains the minimal effect of this mutation on crystal resolution.

**Figure 5:**
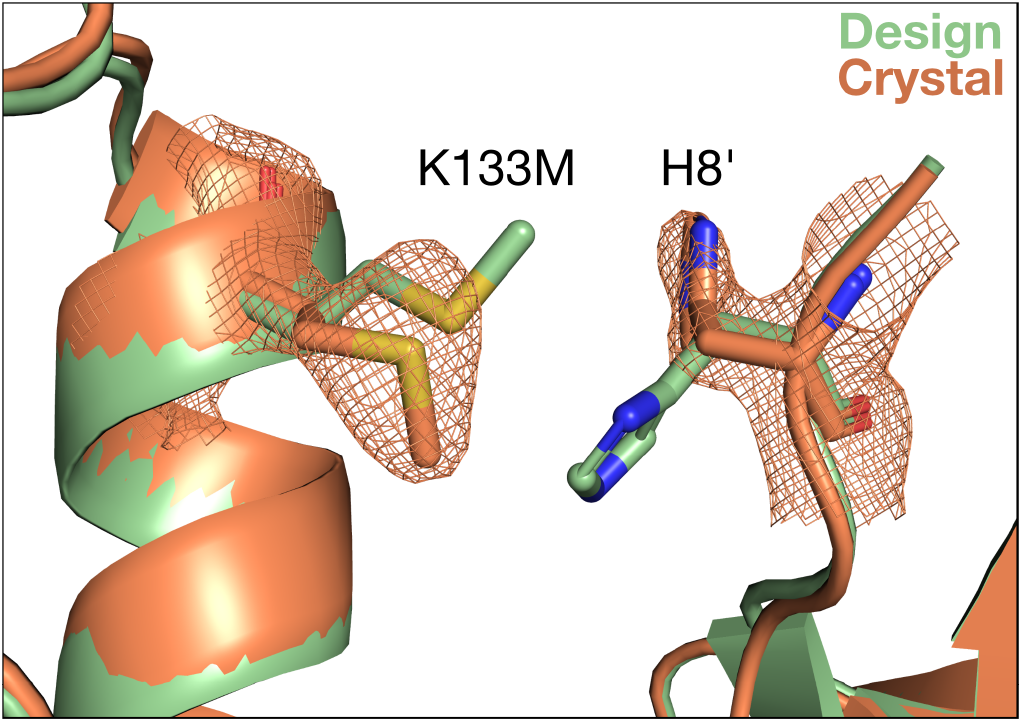
Superposition of the designed (green) and crystallized (orange) K133M structure (PDB ID 6OK9). The 2mFo-DFc map is contoured at 1.5σ and carved within 2 Å of residues 8 and 133. The designed packing interaction (K133M–H8’, where ‘ indicates symmetry mate) does not occur in crystal, instead the side chains occupy alternative rotamers.

#### 3.4.3 K127L and K64R

The variant K127L produced the highest-resolution crystals in our study, though it did so in a manner not predicted by design: it altered the crystal space group. The WT protein crystallizes in the space group P2_1_, and, in this crystal form, K127 forms a salt bridge with the THP molecule bound in the neighboring SNase active site (Figure 6). When this lysine is mutated to leucine, the salt-bridge interaction cannot form, destabilizing the P2_1_ crystal form. Instead, the designed protein crystallizes in P4_1_, a higher symmetry space group, where L127 packs against a neighboring loop by forming backbone interactions with K28 and G29. In this space group, K71 replaces K127 as the interacting partner of the THP in the crystal form, suggesting that the interaction of the substrate phosphate groups with a positively-charged side chain might be useful for SNase crystallization.

**Figure 6:**
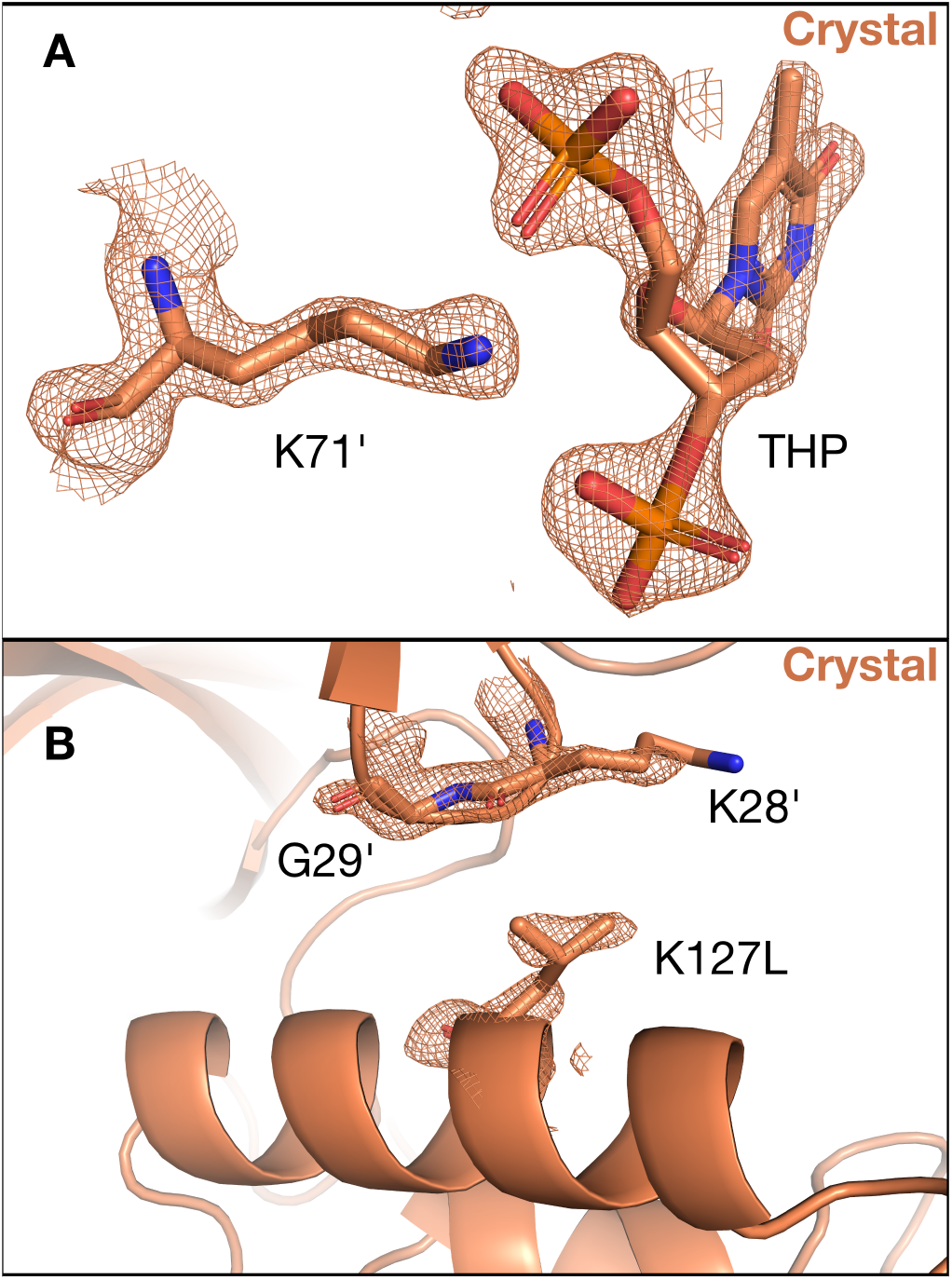
The variant, K127L (PDB ID 6OK8), which yields the highest-resolution crystals, crystallizes in a higher symmetry space group (P 41) than WT (P 21) because it breaks an electrostatic contact central to a crystallographic interface in P 21. (A) In the K127L crystal, the previous K127–THP salt bridge was retained, albeit with a different lysine residue (71). The 2mFo-DFc map is contoured at 1.5σ and carved within 2 Å of residues 71 and the THP molecule. (B) The new crystallographic interface containing L127 is well-resolved in density, and features non-specific side-chain–backbone contacts. The 2mFo-DFc map is contoured at 1.5σ and carved within 2 Å of residues 28, 29 and 127.

In addition to the space group change, the K127L variant had the largest backbone motions of all variants. The motions are in the loop region (residues 114–118) that precedes the α-helix containing L127. They occur when residue K116 shifts from contacting the neighboring molecule in the crystal to make contacts to the bound nucleotide analog instead.

The other substitution that improved resolution in our study, albeit with a sample size of one, was K64R. Although we were unable to solve a crystal structure due to problems with twinning and the presence of ice, we were able to determine from the diffraction data that K64R (like K127L) resulted in a change in space group, going from P2_1_ to the higher symmetry space group P2_1_2_1_2_1_. To gain structural insight as to why this variant and space group might lead to higher resolution, we aligned our K64R model to a different SNase structure crystallized in the same space group (PDB ID: 5KEE) and applied the symmetry operations necessary to generate the neighboring symmetry mates, using unit cell dimensions from the diffraction data. We then used Rosetta’s FastRelax protocol to alleviate any clashes that might have been introduced. We observed two possible hydrogen bonding interactions for R64 that might account for this change in space group: one with the carboxylic acid of E135 of a neighboring symmetry mate and another with the backbone carbonyl of the oxygen of the same residue (Figure 7).

**Figure 7:**
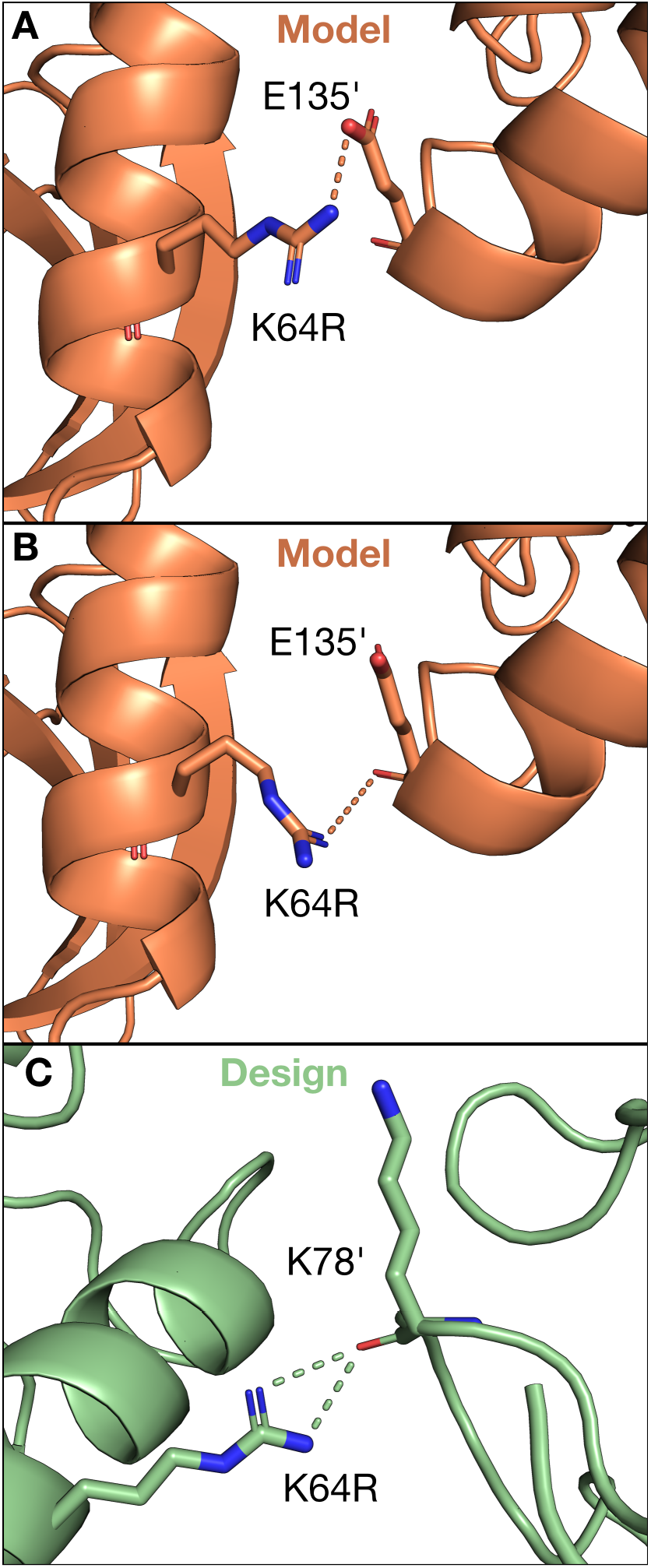
Models (orange) of the possible K64R interactions in the P 2_1_2_1_2_1_ space group show two new possible electrostatic interactions with residue E135 of a neighboring symmetry mate (indicated by the ‘). K64R either interacts with the side-chain or backbone atoms of E135. Residue K64 was not strongly interacting in WT, showing multiple possible rotameric states in electron density and missing density for some atoms (Supplemental Figure 4). Thus, the introduced R64–K78’ interaction in the design (green) was intended to be stabilizing.

#### 3.4.4. Retrospectively: Rosetta Score Recovers Space Group Changes, But Not Resolution

Since we did not include the possibility of space group changes in our design protocol, yet we observed changes for two variants, we retrospectively asked whether Rosetta could recover the correct space group (Figure 8) by modeling and scoring each variant and the WT in each of the three most popular SNase space groups. For four out of six crystals (including WT), we found that Rosetta could correctly predict the space groups. Rosetta failed to predict the correct space group changes for the K127L variant, yielding P2_1_2_1_2_1_ as the lowest scoring space group (when the actual space group was P4_1_), and for the K64R substitution, yielding P2_1_ as the lowest scoring space group (the actual space group was P2_1_2_1_2_1_).

**Figure 8:**
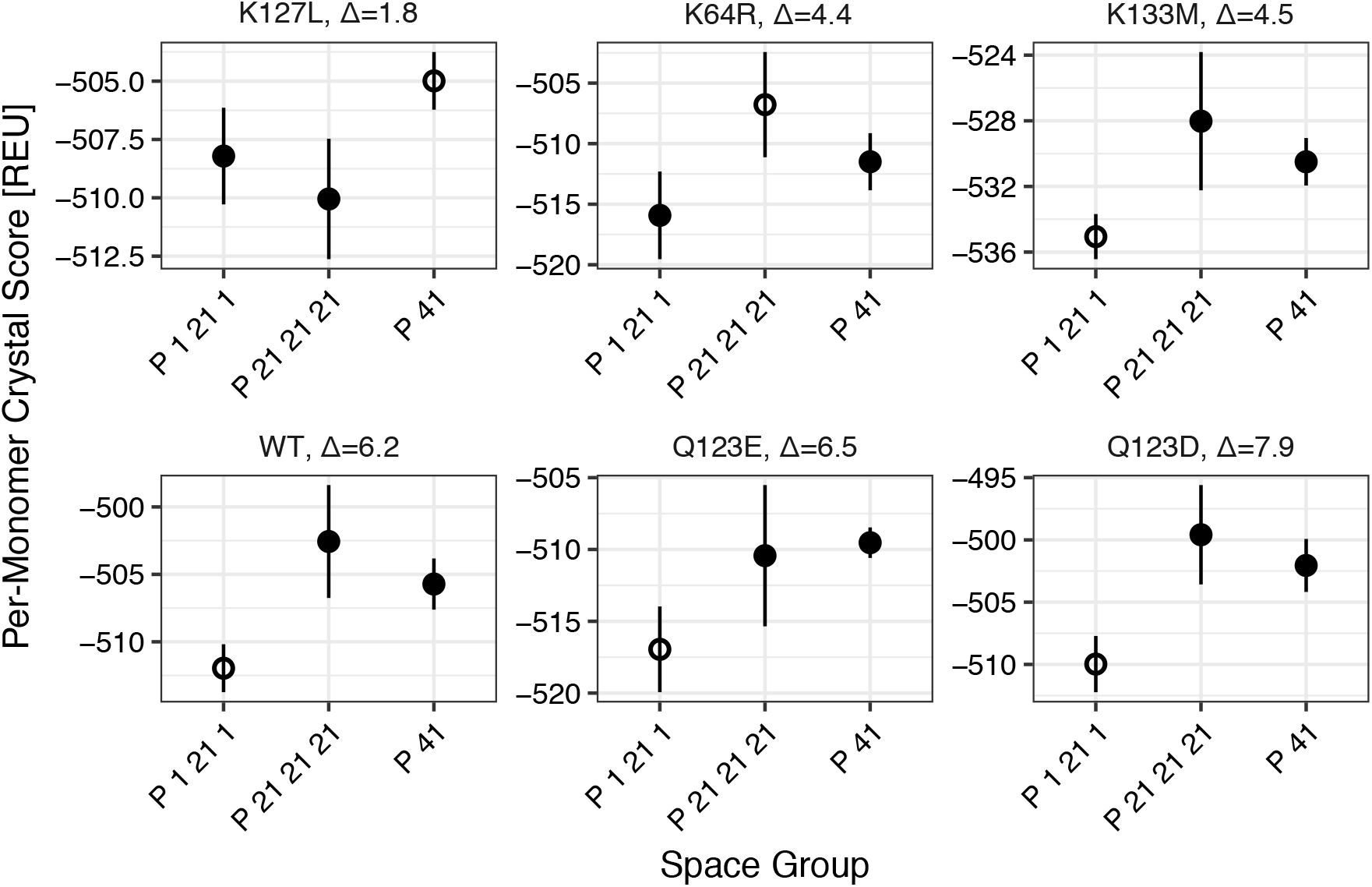
Scores of designs in the three most popular space groups for SNase. The lowest-scoring space group is the experimentally observed space group in four out of six cases. For each design, average scores are shown ± one standard deviation, computed over ten energy-minimized structures in P2_1_, P2_1_2_1_2_1_, and P4_1_. The experimentally observed space group is indicated by an open circle. The designs are ordered by the score difference, Δ, between the lowest scoring and second lowest scoring space group. The Δ value was greater on average for the designs crystallizing in P2_1_ (~6.3 REU vs. ~3.6 REU). For the two designs (K127L and K64R) that did not crystallize in P2_1_, the correct space group could not be identified based on score.

Finally, we asked whether the Rosetta score of the solved crystal structures correlated with the resolution, as we found to be the case for the engineered variants studied by Mizutani *et al.*^17^ (Results 3.1). We analyzed the crystal structures of our designs as previously described. Surprisingly, we found an anti-correlation between score and resolution (Figure 9), although it should be noted that our resolution range only spans ~0.1 Å, whereas ~0.8 Å was spanned by the crystal structures from Mizutani *et al.* (Figure 1).

**Figure 9:**
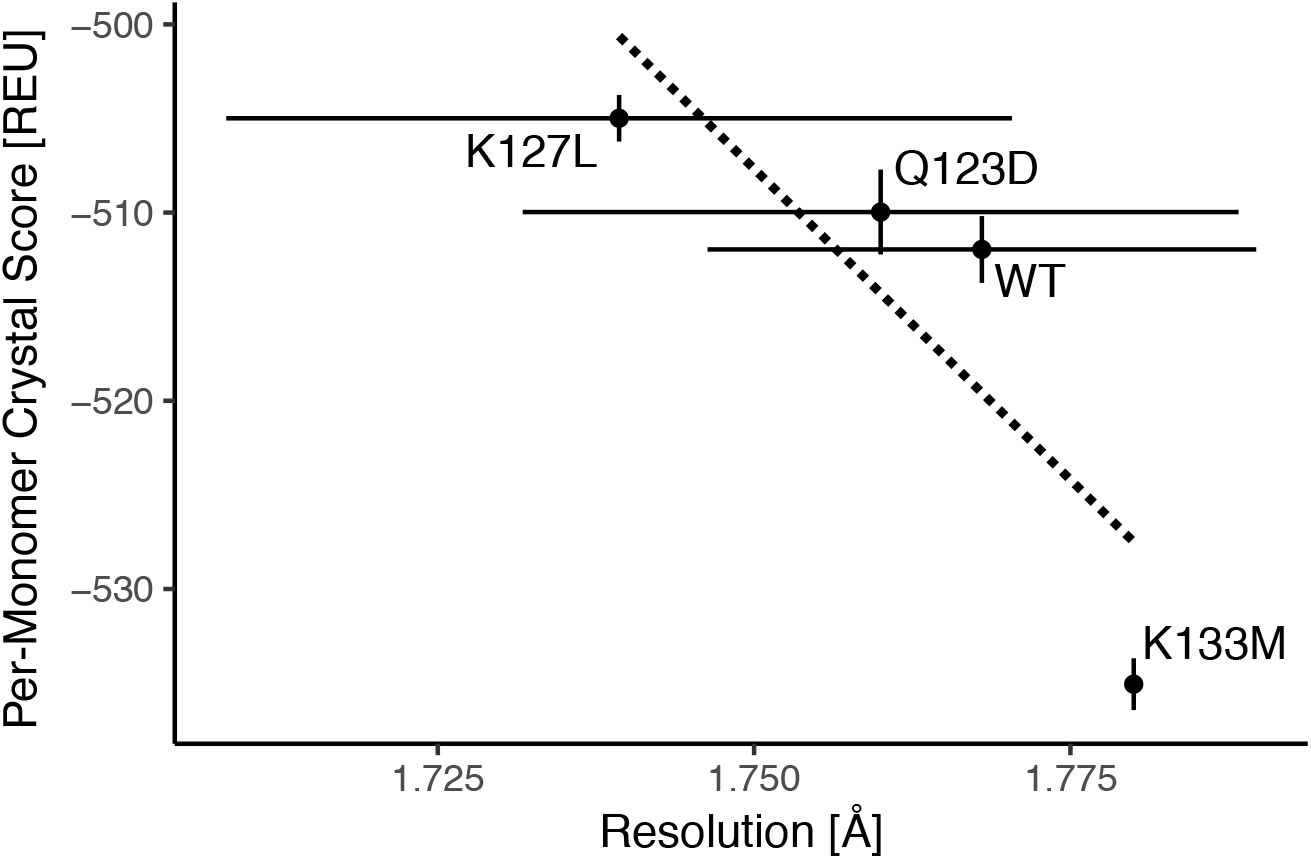
*Ex post facto* analysis by extracting the score from the solved crystal structures of the designs shows an unexpected anti-correlation between resolution and score. Error bars indicate the standard deviation in resolution (from collecting and analyzing multiple diffraction patterns) and score (from ten repeated energy-minimizations).

## 4 Discussion

We attempted to develop and validate a generic computational method for protein crystal contact design to engineer crystals that yield high-resolution structural information. Probing the PDB, we found that Rosetta score correlated with crystal resolution when accounting for common external variables relevant to crystallization such as the decision making to select the highest resolution shell, user handling of the crystal, or crystallization conditions. Using data from an existing study in crystal engineering^17^, we developed a design approach that recapitulated resolution-enhancing mutations at a rate of at least 11.5% (and at best 35.3%). We tested our design approach on a model SNase system, Δ+PHS, and found that our initial approach only resulted in one crystallizable variant out of ten, so we improved our approach by designing on an ensemble of backbones to increase this ratio to four in five. Finally, we solved the crystal structures of several of our designs but unfortunately observed little to no improvement in resolution. *Post-facto* analysis revealed that (1) improvements in resolution came primarily from changes in space group and (2) Rosetta score of designs was not predictive of crystal resolution.

### 4.1 Point Mutations Primarily Affected Side-Chain Interactions and Space Groups

In general, when all variants were compared to both the WT and predicted design structures, the changes in the fold of SNase were undetectable, but there were detectable changes in side-chain interactions at the crystallographic interfaces. First, there were minimal changes in backbone structure, as anticipated for variants that differ by only point mutations at surface residues. The maximum observed root-mean-squared deviation (RMSD) for backbone atoms (N, C, CA, O) between the designed model (or WT, since the backbone was fixed during design) and variant crystal structure was 0.53 Å for the variant K127L, with all other variants having lower backbone RMSD to their respective designed model (or WT, Supplemental Figure 2). Second, all mutations had some unpredicted effects on the interactions at the targeted crystallographic interface. These effects ranged from slight differences in side-chain rotameric states to entirely new interfaces. The smallest number of differences was observed in the K133M variant, where only a few side-chain dihedral angles differed from the designed structure and the interface was not greatly perturbed in general (Figure 5).

The greatest difference we observed was that two variants crystallized in higher symmetry space groups than WT (P2_1_): K64R crystallized in the space group P2_1_2_1_2_1_, and K127L crystallized in the space group P4_1_. Several observed improvements in resolution were seen for this additional symmetry, such as a consistently higher I/σ value (a measure of the information content) over all resolution shells (Supplemental Figure 4). The space group change for K127L was driven by breaking the WT lysine– THP contact across one crystallographic interface (Figure 5), whereas the driver for the space group change of K64R was not definitively determined. The K64R substitution was desired because, in the WT space group (Figure 7), it introduced a putative electrostatic interaction between a terminal amino group of R64 and the carbonyl oxygen of K78 of a symmetry mate,. The mutation did not disrupt any contacts, as K64 was not resolved in the electron density of the WT crystal structure (Supplemental Figure 5); however, this alteration still resulted in a change in space group. Since we were unable to solve a crystal structure for this variant, we resorted to modeling K64R in the new space group. Our models hinted that this space group might be preferred over the WT because R64 can potentially form a hydrogen bond with E135 via both side-chain–side-chain and side-chain–backbone interactions (Figure 7).

### 4.2 The Relationship Between Score and Resolution

Our initial hypothesis was that crystal interface stability, as captured by Rosetta score, would correlate with crystal resolution. We had two reasons that led us to this hypothesis. First, crystal growth occurs when the rate of protein incorporation into a crystal lattice (attachment) is greater than the rate of protein detachment from the crystal. The rate of attachment depends on the flux of molecules to the growing crystal step, the possible interaction area of the step, and the probability of attachment. The rate of detachment depends on the frequency of detachment events and the detachment probability. Point mutations can affect both the detachment and attachment probabilities. The detachment probability in particular is related to interface stability by the Boltzmann factor: *p*_*d*_ = exp (−*E*_*i*_/*kT*)^42^. All other factors being equal, more stable native crystal interfaces will result in lower probabilities of detachment and thus may improve crystal morphology. Second, as interface stability is conferred by favorable molecular interactions and tighter side-chain packing, we reasoned that more stable interfaces would be less dynamic and less mobile, improving the homogeneity of uniform protein positioning within the crystal. Hence, we anticipated that more stable interfaces would result in larger crystals with less mosaicity, which in turn would improve crystal resolution.

An initial analysis comparing Rosetta scoring and crystal resolution for a subset of the PDB revealed no relationship between the two. We reasoned that our analysis was obfuscated by many factors that affected resolution, but could not be captured by score alone (*e.g.* the protein or X-ray source intensity or detector resolution, user handling, etc.). First, we controlled for just the protein by analyzing only structures of SNase in the PDB. We found that controlling for the protein alone was insufficient — there was no trend between resolution and score for this set. However, when we controlled for more factors by analyzing the crystal structures of twenty-two variants of a single protein, with all data gathered by the same individual, using the same process, and with the same equipment, we found, as hypothesized, that lower Rosetta scores correlated with higher resolution (Figure 1). Yet, when we repeated the same analysis for crystals of our model protein (again with all experiments conducted identically by one individual), we found an anti-correlation.

Why is there an inconsistent behavior between score and resolution? From the PDB, it is apparent that higher resolution crystal structures tend to have better protein geometry, *i.e.* fewer improbable side-chain rotamers, fewer outliers for bond lengths, fewer outliers for bond angles, or fewer atomic/steric clashes^43^. It is possible that because Rosetta represents proteins in internal coordinate space (Φ/Ψ), with fixed bond lengths and angles, it cannot rescue inherently poor geometry and thus better geometry contributes to a lower Rosetta score, even after energy minimization. Then, it is possible that the correlation observed between Rosetta score and resolution for the Mizutani set was a manifestation of the protein geometry improvements that come with higher resolution data, while some external factor, unaccounted for by Rosetta score, affected resolution. If resolution does indeed drive score, then for crystals in a narrow range of resolutions, we would not expect to observe a correlation between Rosetta score and resolution, as was the case for the Δ+PHS variants. In fact, MolProbity^44^, a structure validation software, only compares structures within 0.25 Å bins to account for the improvements in protein geometry offered by higher resolution data.

### 4.3 Backrub Improves Design

Over the course of this study, we attempted both fixed-backbone design on the WT backbone and fixed-backbone design on a perturbed ensemble of 200 models generated from the WT backbone. To generate the perturbed ensemble, we used an approach known as Backrub^26^ that slightly alters the direction of the Cα–Cβ vector to expose the side chain to a new environment while minimally altering the backbone. We found that variants designed using an ensemble of backbones crystallized at a higher rate (4/7) than variants design using the single WT backbone (1/11). We reason that this is because Backrub-generated ensembles capture local backbone fluctuations, whereas fixed-backbone models do not, resulting in a better estimate of point mutation effects, including to interface energy^27^. In general, it is known that proteins are dynamic and are readily capable of incorporating point mutations, especially at protein surface positions^45^, so a fixed-backbone approximation is not sufficient. Hence, we observed an increase in crystallization success rate when we designed on an ensemble of backbones and selected designs scoring well across multiple backbones for experimental characterization.

### 4.4 What We Couldn’t Predict and Why

Of the five designed proteins, only K133M resulted in a crystal structure similar to the design. The designs Q123D and Q123E resulted in the introduction of E/D–K electrostatic interactions, but with a neighboring symmetry mate instead of the one targeted by the design. For these designs, it is not clear how to improve the design algorithm.

For the K127L and K64R designs, we observed unpredictable changes in space group. In retrospect, the K127L design should not have scored well in Rosetta, as the K127 amino group clearly makes electrostatic contacts with the phosphate groups of the THP molecule bound by the neighboring symmetry mate. Despite breaking the lysine–THP interaction, the Rosetta score was lower for the variant than the WT protein (in the WT space group), indicating that Rosetta does not correctly weigh the strength of this electrostatic interaction. One possible solution to overcome this issue in the future would be to bias the Rosetta score by the WT electron density, such that eliminating a clearly present interaction is strongly penalized whereas designing residues that are not well-resolved is favored.

One possible reason for the failure to improve resolution is that Rosetta is not yet finely tuned for the types of atomic interactions we tried to create. Rosetta was first developed to study protein folding in the context of small, globular domains, before being applied to the inverse challenge, design^46^. Over the years Rosetta has performed best when redesigning protein cores and tightly-packed interfaces^47–50^. Our objective here is one of the first attempts to design a loosely-packed interface with a significant amount of water involved. Future work to improve design of solvated interfaces might include explicitly analyzing water interactions the interface either by flooding, as was recently successfully used to dock interfacial waters^51^, or the recently developed HBNet method in Rosetta, which has been used to design hydrogen bonding networks *de novo*^52^. Multi-state design^53^ might also be necessary to prevent undesired changes in space group.

### 4.5 Were Our Model Protein Crystals Already Optimal?

Initial analyses showed that diffraction patterns collected for the WT control in this study had an average high-resolution limit of 1.77 Å (Figure 3), which agrees with the resolution (1.8 Å) of the PDB-deposited crystal structure of Δ+PHS (3BDC). This value falls firmly in the middle of the distribution of all SNase crystal resolutions (Figure 1), with 1.35 Å and 2.5 Å being the highest and lowest observed resolutions, respectively. Separately, Mizutani *et al.* observed changes from +0.2 to −0.6 Å in their study of the effects of point mutations on diphthine synthase crystals^17^. Based on these prior observations, we expected to observe changes in resolution of ±0.5 Å; however, we instead found that designs spanned a narrow range of 1.67–1.85 Å or ~1.77±0.1 Å. Nonetheless, the variance in resolution within crystals of the same variant compared favorably between our study and that by Mizutani *et al.* We observed ranges of ~0.1 Å, while Mizutani *et al.* reported a 95% confidence interval estimate of ±0.05 Å for WT diphthine synthase, analyzing the diffraction data from 13 crystals^17^.

One possible explanation for our designs’ minimal improvement in resolution is that our choice in model protein, Δ+PHS, was already optimized for forming high-quality crystals. We selected Δ+PHS as a model system for its high stability (11.8 kcal/mol)^29^, high yield (over 60 mg protein per 1 L of cell culture), and crystallizability (over 300 crystal structures have been deposited in the PDB). We selected for these features so that our model protein would readily incorporate point mutations and so that the corresponding designs would likely express in high quantities and readily crystallize. However, these features also likely pre-selected for a protein that is optimal for crystallization, one for which a majority of point mutations might not be able to yield significant improvements in resolution. Future work might then focus on a protein that is less engineered and may have more room for improvement.

## 6 Author Contributions

JRJ, ACR, JMB, BGME and JJG designed the research. JRJ and ACR performed the research. JRJ, ACR, JMB, BGME, and JJG analyzed the data. JRJ, ACR, JMB, BGME, and JJG wrote the paper.

## 8 Acknowledgments

The authors would like to acknowledge Jesse B. Yoder for helpful discussion and Michael L. Love for help with instrumentation. The super-computing resources in this study have been provided in part by the Maryland Advanced Research Computing Center. Diffraction data was collected at the X-ray laboratory of the Department of Biophysics and Biophysical Chemistry, School of Medicine, Johns Hopkins University.

