## Supplemental Figures for "Toward computational design of protein crystals with improved resolution"

**Supplemental Figure 1**: **The ability for Rosetta to correctly forward design resolution-enhancing mutations on diphthine synthase depends on the degrees of freedom sampled during the simulation.** Each plot here shows the same data as Figure 2 for a particular design strategy. The x-axis indicates the variant resolution and the y-axis shows the difference in energy with respect to the WT (hence the dashed lines represent the WT values for both). Standard deviations are calculated across 10 repeated design simulations. Point color indicates either a correct (*i.e. a lower score than WT and corresponding higher resolution*, blue) or incorrect prediction (red). The strategies are fully detailed in the Results section. Neither the strategy with the most (*Crystal Docking*, #6 in Results) nor the fewest (*Repack*, #1) degrees of freedom sampled was particularly successful. In fact, minimizing on side-chain (*Repack and SC Min*, #2) or rigid-body degrees (*Repack and SC, Jump Min*, #4) of freedom was not sufficient. Rather, successful strategies featured backbone minimization after the point mutation was introduced (*Repack and SC, BB Min*, #3, or *Repack and SC, BB, Jump Min*, #5).

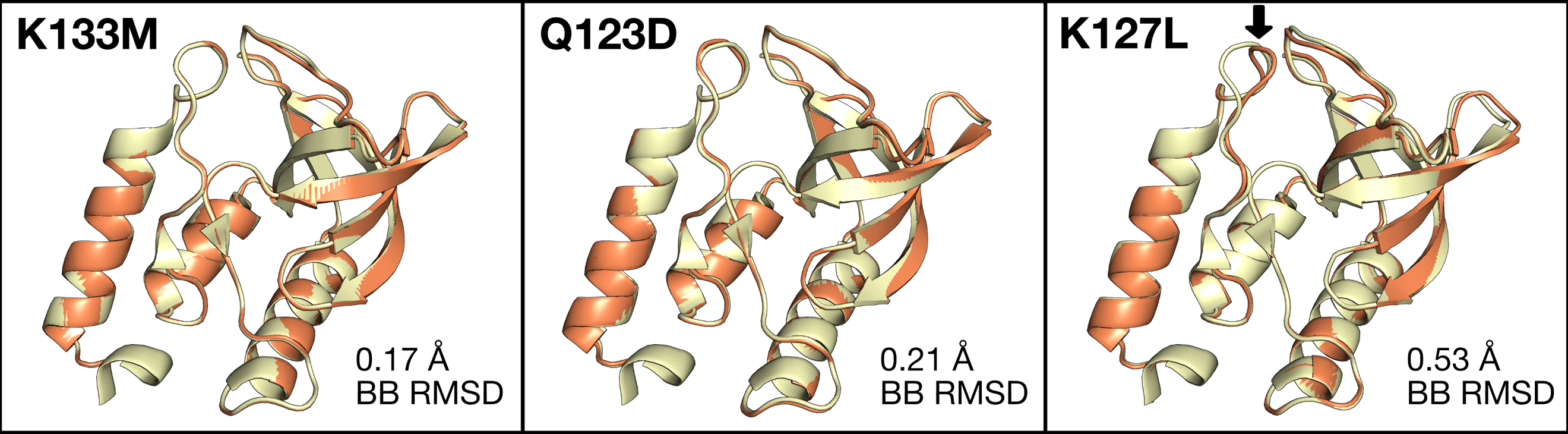

**Supplemental Figure 2**: **Alignments between design (orange) and crystal structure (yellow) backbones.** There are minimal differences, with the greatest backbone RMSD being 0.53 Å for the K127L variant.

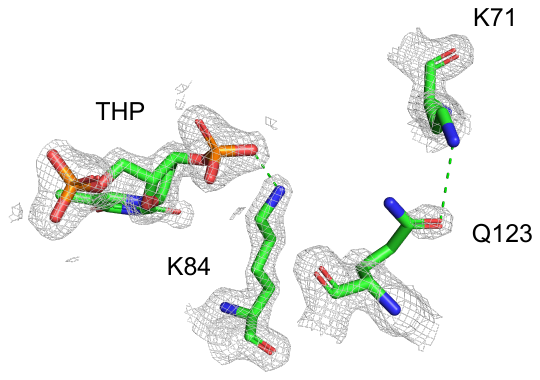

**Supplemental Figure 3: WT interactions between THP and K84, and Q123 and K71.** Coordinates for the WT come from PDB ID 3BDC. The 2mFo-DFc map was downloaded from the Uppsala Electron Density Server. The map is contoured at 1.5σ and carved within 2 Å of residues 71, 84, 123, and the THP. Distances below 3.5 Å are highlighted by green dashed lines.

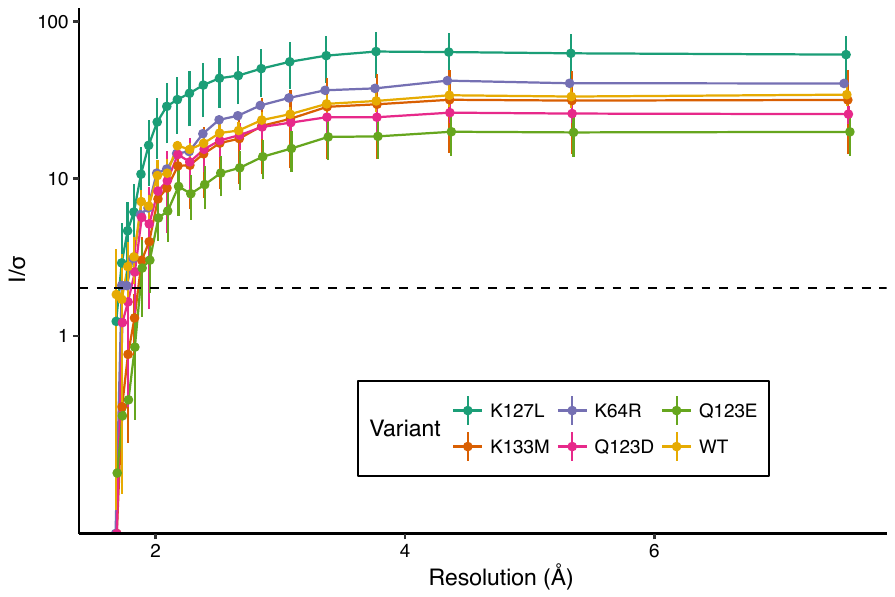

**Supplemental Figure 4**: **Average I/σ values ± one standard deviation for each resolution shell for each variant.** Averages and standard deviations are calculated over all protein crystals for which diffraction data could be collected.

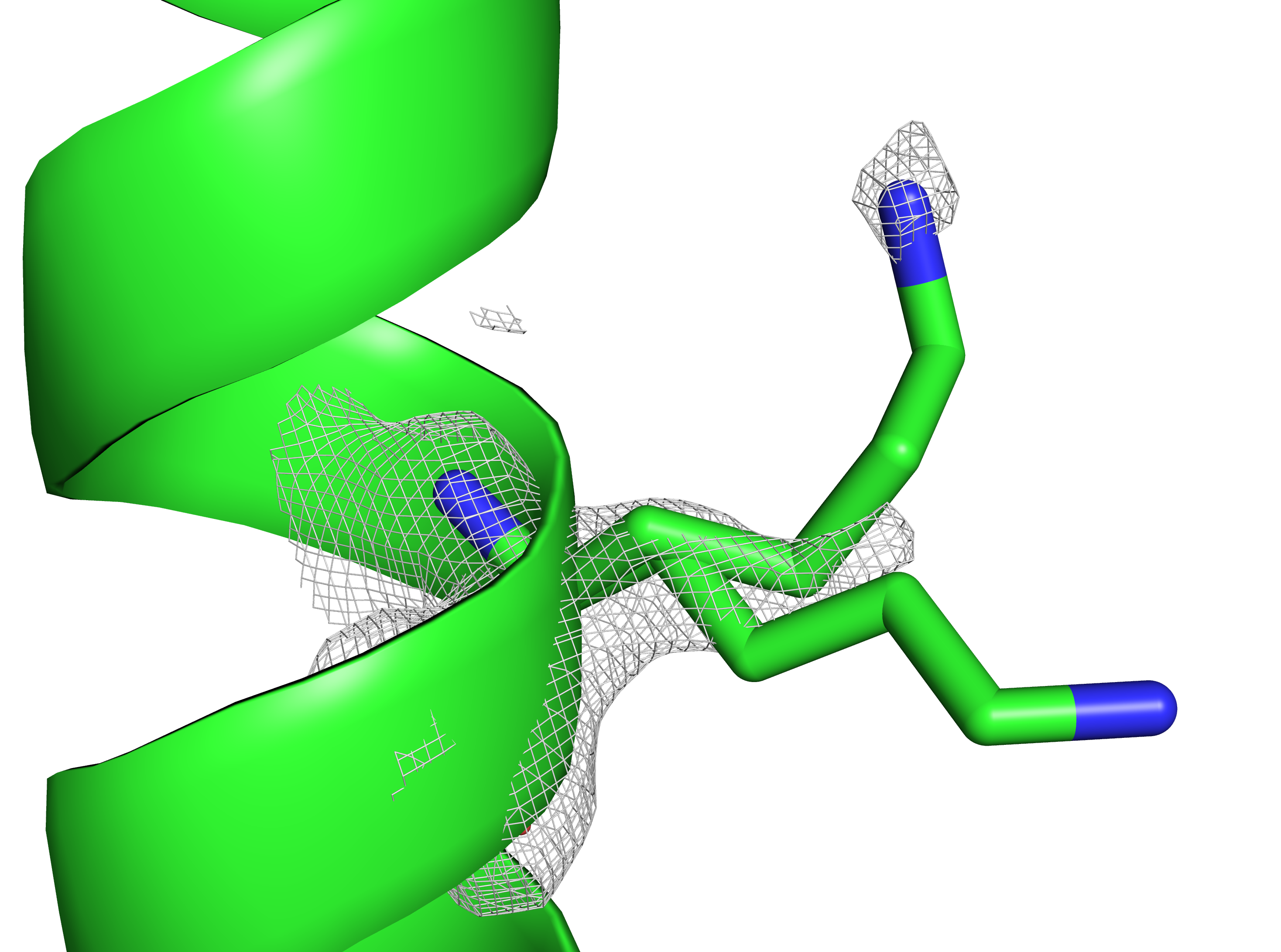

**Supplemental Figure 5: WT density of K64 is missing for some atoms and shows multiple rotameric states.** Coordinates for the WT come from PDB ID 3BDC. The 2mFo-DFc map was downloaded from the Uppsala Electron Density Server. The map is contoured at 1.5σ and carved within 2 Å of residue 64.
