## Supplemental Tables for "Toward computational design of protein crystals with improved resolution"

**Supplementary Table 1**: **Crystallization conditions and I/σ and completeness values for the highest-resolution shell observed.**

| Variant | % MPD | pH | Ca2+ ratio | pdTp ratio | Space Group | Resolution | I/σ | Completeness (%) |
| --- | --- | --- | --- | --- | --- | --- | --- | --- |
| Δ+PHS K127L | 40 | 9 | 3 | 2 | P41 | 1.67 | 3.36 | 9 |
| Δ+PHS K127L | 40 | 9 | 3 | 2 | P41 | 1.77 | 4.74 | 48.8 |
| Δ+PHS K127L | 40 | 8 | 3 | 2 | P41 | 1.69 | 2.37 | 9.5 |
| Δ+PHS K127L | 42 | 9 | 3 | 2 | P41 | 1.77 | 2.23 | 46.3 |
| Δ+PHS K127L | 42 | 9 | 3 | 2 | P41 | 1.69 | 5.1 | 9 |
| Δ+PHS K127L | 42 | 9 | 3 | 2 | P41 | 1.68 | 3.59 | 7.7 |
| Δ+PHS K127L | 42 | 8 | 3 | 2 | P41 | 1.73 | 4.78 | 25 |
| Δ+PHS K127L | 44 | 9 | 3 | 2 | P41 | 1.83 | 2.72 | 79.6 |
| Δ+PHS K127L | 44 | 8 | 3 | 2 | P41 | 1.73 | 4.57 | 23.2 |
| Δ+PHS K127L | 44 | 8 | 3 | 2 | P41 | 1.78 | 2.48 | 44 |
| Δ+PHS K127L | 46 | 9 | 3 | 2 | P41 | 1.68 | 2.98 | 7.7 |
| Δ+PHS K127L | 46 | 9 | 3 | 2 | P41 | 1.73 | 6.77 | 26.4 |
| Δ+PHS K127L | 46 | 8 | 3 | 2 | P41 | 1.68 | 2.29 | 8.7 |
| Δ+PHS K127L | 46 | 8 | 3 | 2 | P41 | 1.83 | 3.05 | 69.5 |
| Δ+PHS K127L | 46 | 8 | 3 | 2 | P41 | 1.78 | 2.34 | 46.4 |
| Δ+PHS WT | 21 | 6 | 3 | 2 | P21 | 1.73 | 3.35 | 11.4 |
| Δ+PHS WT | 18 | 6 | 3 | 2 | P21 | 1.77 | 3.33 | 33 |
| Δ+PHS WT | 18 | 6 | 3 | 2 | P21 | 1.78 | 2.74 | 25.9 |
| Δ+PHS WT | 18 | 6 | 3 | 2 | P21 | 1.74 | 3.6 | 12.1 |
| Δ+PHS WT | 18 | 6 | 3 | 2 | P21 | 1.83 | 2.08 | 45.4 |
| Δ+PHS Q123E | 18 | 6 | 2 | 1 | P21 | 1.89 | 2.79 | 68.8 |
| Δ+PHS Q123E | 20 | 6 | 2 | 1 | P21 | 1.94 | 2.62 | 66.2 |
| Δ+PHS Q123E | 20 | 6 | 2 | 1 | P21 | 1.89 | 3.61 | 56.5 |
| Δ+PHS Q123E | 20 | 6 | 2 | 1 | P21 | 1.84 | 2.37 | 24.6 |
| Δ+PHS Q123E | 18 | 6 | 2 | 1 | P21 | 1.89 | 5.14 | 60.2 |
| Δ+PHS Q123E | 18 | 6 | 2 | 1 | P21 | 1.88 | 2.64 | 67 |
| Δ+PHS Q123E | 18 | 6 | 2 | 1 | P21 | 1.96 | 2.37 | 87.1 |
| Δ+PHS K133M | 20 | 6 | 2 | 1 | P21 | 1.84 | 2.16 | 43.7 |
| Δ+PHS K133M | 18 | 6 | 2 | 1 | P21 | 1.89 | 2.48 | 67.4 |
| Δ+PHS K133M | 18 | 6 | 2 | 1 | P21 | 1.9 | 3.98 | 81.6 |
| Δ+PHS K133M | 18 | 6 | 2 | 1 | P21 | 1.79 | 2.62 | 26.6 |
| Δ+PHS K133M | 22 | 6 | 2 | 1 | P21 | 1.89 | 3.58 | 74.3 |
| Δ+PHS K64R | 18 | 6 | 3 | 2 | P212121 | 1.73 | 2.13 | 22.2 |
| Δ+PHS Q123D | 20 | 6 | 2 | 1 | P21 | 1.95 | 2.39 | 87.9 |
| Δ+PHS Q123D | 20 | 6 | 2 | 1 | P21 | 1.78 | 2.61 | 25.9 |

**Supplemental Table 2**: **Selected statistics for deposited crystal structures.**

|  | **K127L (6OK8)** | **K133M (6OK9)** | **Q123D (6OKA)** |
| --- | --- | --- | --- |
| **Wavelength** | 1.54 | 1.54 | 1.54 |
| **Resolution range** | 38.26–1.8 (1.865–1.8) | 32.21–1.901 (1.968–1.901) | 32.26–1.86 (1.927–1.86) |
| **Space group** | P 41 | P 1 21 1 | P 1 21 1 |
| **Unit cell** | 48.097 48.097 63.122  90 90 90 | 30.927 60.473 38.105  90 93.017 90 | 30.853 60.715 38.119  90 92.648 90 |
| **Total reflections** | 72360 (1421) | 33219 (1633) | 34560 (1407) |
| **Unique reflections** | 12307 (715) | 10702 (874) | 11133 (680) |
| **Multiplicity** | 5.9 (2.0) | 3.1 (1.9) | 3.1 (2.1) |
| **Completeness (%)** | 91.93 (53.32) | 96.32 (79.71) | 93.59 (56.06) |
| **Mean I/sigma(I)** | 35.73 (5.59) | 25.62 (3.92) | 18.37 (5.10) |
| **Wilson B-factor** | 22.28 | 29.02 | 28.23 |
| **R-merge** | 0.02957 (0.1052) | 0.02397 (0.1765) | 0.03833 (0.1442) |
| **R-meas** | 0.03219 (0.1381) | 0.02873 (0.2304) | 0.04588 (0.1879) |
| **R-pim** | 0.01254 (0.08851) | 0.01565 (0.146) | 0.02494 (0.119) |
| **CC1/2** | 1 (0.984) | 0.999 (0.983) | 0.998 (0.978) |
| **CC*** | 1 (0.996) | 1 (0.996) | 1 (0.994) |
| **Reflections used in refinement** | 12308 (715) | 10706 (868) | 11125 (680) |
| **Reflections used for R-free** | 619 (38) | 538 (44) | 562 (38) |
| **R-work** | 0.1950 (0.2892) | 0.2199 (0.3940) | 0.1986 (0.3391) |
| **R-free** | 0.2331 (0.4364) | 0.2635 (0.4842) | 0.2532 (0.4043) |
| **CC(work)** | 0.961 (0.826) | 0.956 (0.785) | 0.935 (0.475) |
| **CC(free)** | 0.947 (0.871) | 0.953 (0.818) | 0.941 (0.205) |
| **Number of non-hydrogen atoms** | 1147 | 1058 | 1102 |
| **macromolecules** | 1032 | 993 | 999 |
| **ligands** | 26 | 26 | 26 |
| **solvent** | 89 | 39 | 77 |
| **Protein residues** | 129 | 129 | 129 |
| **RMS(bonds)** | 0.022 | 0.025 | 0.022 |
| **RMS(angles)** | 2.03 | 1.63 | 1.73 |
| **Ramachandran favored (%)** | 92.91 | 94.49 | 93.70 |
| **Ramachandran allowed (%)** | 4.72 | 4.72 | 5.51 |
| **Ramachandran outliers (%)** | 2.36 | 0.79 | 0.79 |
| **Rotamer outliers (%)** | 3.74 | 3.09 | 1.04 |
| **Clashscore** | 1.41 | 2.52 | 3.01 |
| **Average B-factor** | 25.95 | 38.52 | 31.71 |
| **macromolecules** | 25.29 | 38.54 | 31.51 |
| **ligands** | 34.24 | 39.44 | 28.08 |
| **solvent** | 31.23 | 37.34 | 35.60 |
